# Design of a radial multi-offset detection pattern for *in vivo* phase contrast imaging of the retinal ganglion cells in humans

**DOI:** 10.1101/2020.12.08.416826

**Authors:** Elena Gofas-Salas, Yuhua Rui, Pedro Mecê, Min Zhang, Valerie C. Snyder, Kari V. Vienola, Daniel Lee, José-Alain Sahel, Kate Grieve, Ethan A. Rossi

## Abstract

Previous work has shown that multi-offset detection in adaptive optics scanning laser ophthalmoscopy (AOSLO) can be used to image retinal ganglion cells (RGCs) in monkeys and humans. However, though images of RGCs in anesthetized monkeys with high light levels produced high contrast images of RGCs, images from humans failed to reach the same contrast due to several drawbacks in the previous dual-wavelength multi-offset approach. Our aim here was to design and build a multi-offset detection pattern for humans at safe light levels that could reveal the retinal ganglion cell layer neurons with a contrast, robustness and acquisition time approaching results only previously obtained in monkeys. Here, we present a new imaging system using only one light source, compared to the previous dual-wavelength used on monkeys. Our single-wavelength solution allows for increased light power and eliminates problematic chromatic aberrations. Then, we demonstrate that a radial multi-offset detection pattern with an offset distance of 8-10 Airy Disk Diameter (ADD) is optimal to detect photons multiply scattered in all directions from RGCs thereby enhancing their contrast. This new setup and image processing pipeline led to improved imaging of retinal ganglion cells using multi-offset imaging in AOSLO.

## 1. Introduction

The adaptive optics scanning light ophthalmoscope (AOSLO) has been continuously improved over the past 20 year to add new capabilities permitting the detection of different classes of retinal cells. These modifications are usually applied to the illumination and detection path of the system to achieve different imaging modalities, often adapting techniques first applied in microscopy. For example, individual retinal pigmented epithelial cells were detected in AOSLO by adding a fluorescence detection channel with appropriate excitation wavelengths and emission filters [1, 2]. Another exquisite example of a microscopic technique applied for retinal imaging is phase contrast. Transparent or low contrast retinal structures such as photoreceptors inner segments and capillaries were revealed by implementing phase contrast imaging techniques that are sensitive to variation of the refractive index rather than simply absorption or reflectivity [3]. Phase contrast imaging in the retina using AOSLO can be achieved using a range of modalities based on off-axis detection [4]. The simplest configuration shifts the confocal aperture from its on-axis position to reject the single backscattered photons from highly reflective layers and detect the multiply scattered light that is forward scattered by retinal structures with low reflectivity. This configuration, called offset-aperture AOSLO [5], reveals the wall structure of capillaries with exquisite detail. Another related technique is non-confocal split detection AOSLO. Here the light is split into three channels with the central (confocal) portion directed to a first detector, while the light outside the confocal region is split with a mirror and sent to two additional detectors. Images from the two detectors are combined by computing the normalized difference between the images divided by their sum and reveals the photoreceptor inner segments even when the outer segments are degenerated [6].

It has been shown that the scanning beam is deviated in different directions away from the optical axis when going through refractive index gradients within and between single cells, with the amount of steered photons depending on the refractive index gradient magnitude and sign [3, 7]. Thus these previously described methods were only capturing part of the translucent cellular information leaving an important number of deviated photons carrying the phase signal undetected. Rossi *et al*. [8] combined the principles of these two non-confocal techniques and developed multi-offset imaging by laterally displacing a pinhole around the optical axis and computing linear combinations between the images obtained at multiple positions across the retinal conjugate focal plane in AOSLO. A confocal image was obtained simultaneously using a different wavelength to permit eye tracking and registration of the images across the different aperture positions that were acquired sequentially. With this off-axis configuration Rossi *et al*. showed the first high resolution images of ganglion cells *in vivo* in monkeys and humans, albeit with lower contrast and signal-to-noise ratio (SNR) in the human.

Achieving *in vivo* imaging of retinal ganglion cells (RGCs) in humans was more challenging than in the monkey due to four main limitations of the imaging system: 1) This dual-wavelength implementation introduced chromatic aberration between the different wavelengths used for multi-offset and confocal imaging, with transverse chromatic aberration causing a lateral displacement of the images and longitudinal chromatic aberration causing them to focus at different positions axially within the retina. This introduces the potential for error when using the confocal image to track the motion of the eye and applying that motion to the offset channels. 2) The dual-wavelength approach also limited the amount of light power for multi-offset imaging that can be sent to the eye (a substantial amount of the light budget was spent on the visible wavelength confocal channel). In addition, lower light levels were used for imaging the humans compared to the monkeys, where monkey imaging was done using light levels much higher (see [8] for details on the light levels used) than those that can be permitted in humans due to safety constraints, resulting in images with substantially lower SNR. 3) Several offsets were used sequentially around the optical axis, rendering the acquisition quite long and therefore fastidious for subjects (affecting the image registration as eye motion increases and adaptive optics performance can decrease due to the dry eyes). 4) All images from different offsets were combined and the best combinations among all images was manually identified based on visual criterion leading to long post-processing and analysis times.

Here, we solve all these drawbacks: We show the design and implementation of a radial single-wavelength multi-offset detection configuration for AOSLO that addresses the limitations for human RGC imaging in multi-offset AOSLO caused by 1) chromatic aberrations and 2) low light levels from our previous dual-wavelength AOSLO. 3) We also evaluate the optimal offset detection configuration with respect to pattern and distance of the pinhole with respect to the optical axis. We show that a radial pattern with 8 offset positions is enough therefore decreasing the acquisition time. Moreover, we further improve the phase contrast of non-confocal AOSLO by decreasing the Airy Disk Diameter (ADD) at the detection plane. 4) We use the SMART method to combine the offset images and generate an image with the highest contrast possible. Finally, we show here that this new system, combined with our improved registration algorithm [9] is capable of robust and repeatable *in vivo* imaging of RGCs in humans.

## 2. Methods

### 2.1. Experimental Setup

#### AOSLO system

Experiments were carried out on the Pittsburgh AOSLO described in detail elsewhere [10]. In summary, this AOSLO system has four illumination sources and four detection channels. Using custom electronics, a maximum of three channels can be recorded simultaneously. This system has multiple modes of imaging, including confocal imaging, near infrared autofluorescence imaging, multi-offset imaging, and visible light imaging. Though we have previously described the entire system and the results from the near infrared autofluorescence and confocal channels [10], this is the first time the multi-offset sub-system is used and fully described. This new optical setup differs substantially from the two AOSLOs that were used previously for multi-offset imaging in humans and monkeys in Rochester [8]. Suppl. Table 1 summarizes and compares the most relevant differences between our new AOSLO and those used in our previous work [8]. The AOSLO we used here also has several features that are designed to facilitate imaging of human participants. Firstly, a custom-made chin and forehead rest with numerous adjustments permits stabilization of the head and a pupil camera enables fast alignment of the position of the eye/head in conjunction with the forehead/chin apparatus being mounted on a motorized translation stage. Secondly, a combination of fixation target and steering mirror is used to help expand the area that can be imaged and at the same time avoid the need for the participant to fixate too far away from the center which may sometimes cause fatigue. The fixation target is projected onto the retina for the participant to follow, allowing fixation-guided imaging of different locations of the retina, while a steering mirror system permits the experimenter fine control of the position of the imaging beam, and positioning to retinal locations beyond those reachable using the fixation target alone.

#### Optical design of Multi-offset detection path

The retina was illuminated with a super-luminescent diode (MS-795-GI15, Superlum, Dublin, Ireland) centered at 795 nm (FWHM=15nm) that is scanned in a raster fashion at frame rate of 30 Hz to across an 1.5° x 1.5° field of view. Light backscattered from the retina is directed towards the detection channel using a dichroic mirror and then focused with an achromatic doublet onto a custom-made mask as shown in the schematic drawing in Fig. 1 (a). The ADD at this plane and confocal PMT plane is 27 μm and remains this value throughout the study. This mask is composed of a fused silica substrate upon which gold was deposited to form a mirror of elliptical shape which act as circular pinhole mirror of 1 ADD when the mask is placed at 45 degree with respect to the incoming beam. As shown in Fig.1, the pinhole mirror reflects the central portion of the PSF to the confocal channel while the remainder of the light distribution (outer portion) is transmitted and then focused with a pair of lenses to a second retinal conjugate focal plane for offset imaging. This second PMT had a lens tube that allowed us to change pinhole according to the imaging protocol and was placed on a motorized xyz-stages (Thorlabs, Inc. RB13 with Z812B servo motor actuators) allowing precise control of the pinhole location.

**Fig. 1.**
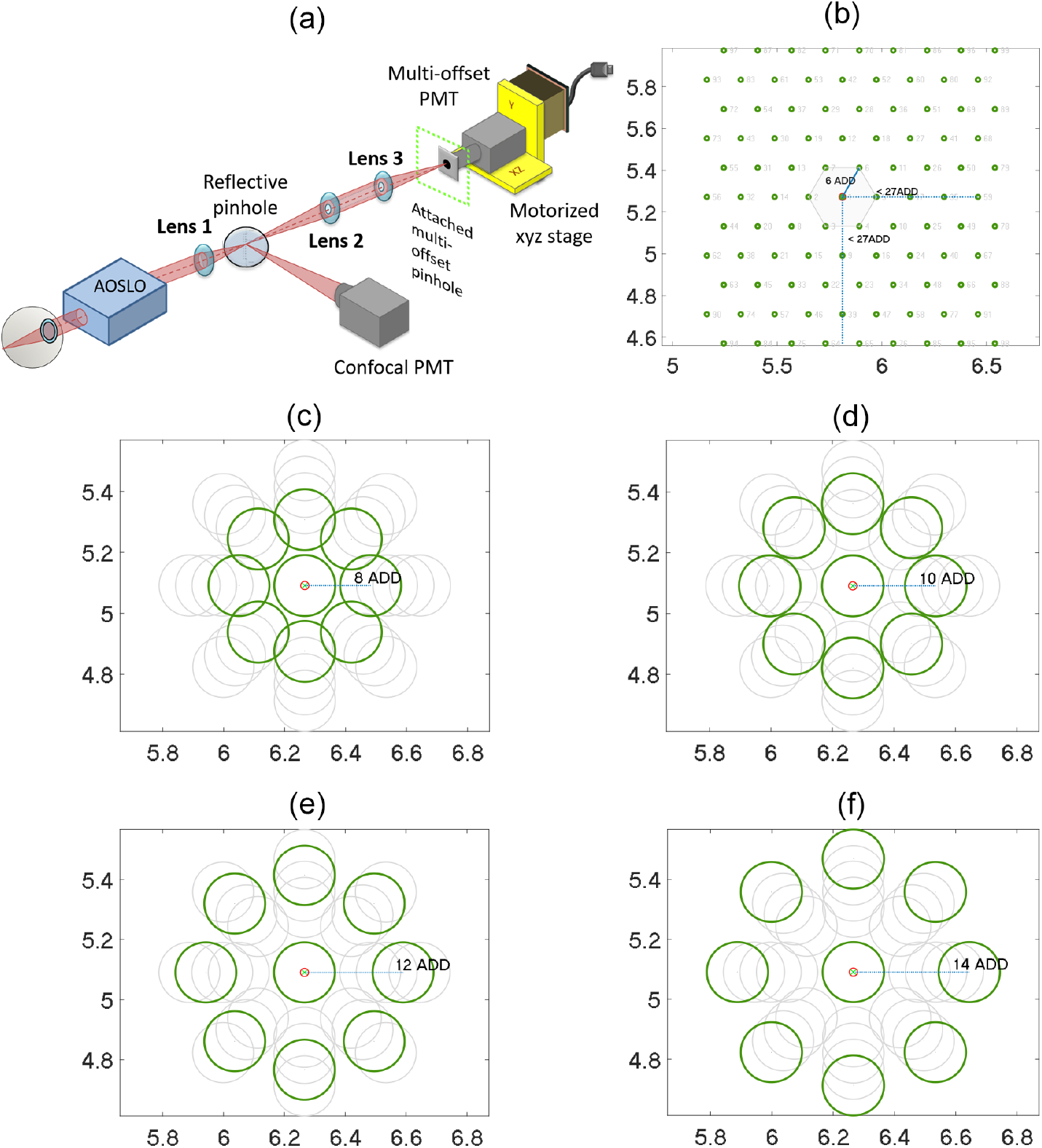
Experimental set-up. (a) Schematic drawing of the single-wavelength multi-offset detection path in the Pittsburgh AOSLO [10]. After passing through the AOSLO and into this detection channel, a first lens (*f*_1_ = 100 mm) focused the light onto a reflective pinhole of 1 ADD size to direct the confocal portion of the light to the confocal PMT. Light outside the confocal area was transmitted and then relayed to a second retinal conjugate focal plane for detection with the multi-offset PMT. Lens 2 has a focal length of *f*_2_ = 100 mm. Lens 3 has a focal length of *f*_3_ = 100 mm or 50 mm depending on the setting (see Table 1). The multi-offset PMT has another pinhole attached and is mounted on a motorized translation stage for precise positioning across the retinal conjugate focal plane. (b) Detection pattern followed by the multi-offset stage for the ‘multi-offset scattering study’ (see Table 1) using a small detection pinhole of less than 1 ADD. (c-f) Detection patterns followed by the multi-offset stage for ‘multi-offset retinal structure imaging study’ (see Table 1) using a larger pinhole of 8 ADD at offset distances {8, 10, 12, 14} ADD respectively. This schematic was inspired by Fig. 1 of Scoles et. al [6].

#### Multi-offset detection pattern

As stated above, the confocal PMT stayed fixed during imaging while the offset channel PMT moved across the xy plane (perpendicular to the optical axis) recording images at different positions. Multi-offset detection pattern refers to the offset PMT positions during the imaging session with reference to the center position. While the confocal PMT plane remained the same throughout this study (27 μm), the ADD at the offset PMT plane was tested with two different values, 27 μm and 13.5 μm; this was achieved by changing the last lens in front of the multi-offset PMT (Lens 3 in Fig. 1(a)). It should be noted that when the ADD changed (from 27 μm to 13.5 μm) at the offset PMT plane, the pinhole aperture and offset distance changed accordingly in the unit of micrometers but remains the same scale in the unit of ADD. Table 1 lists the 5 detection patterns we used in our study.

**Table 1.**
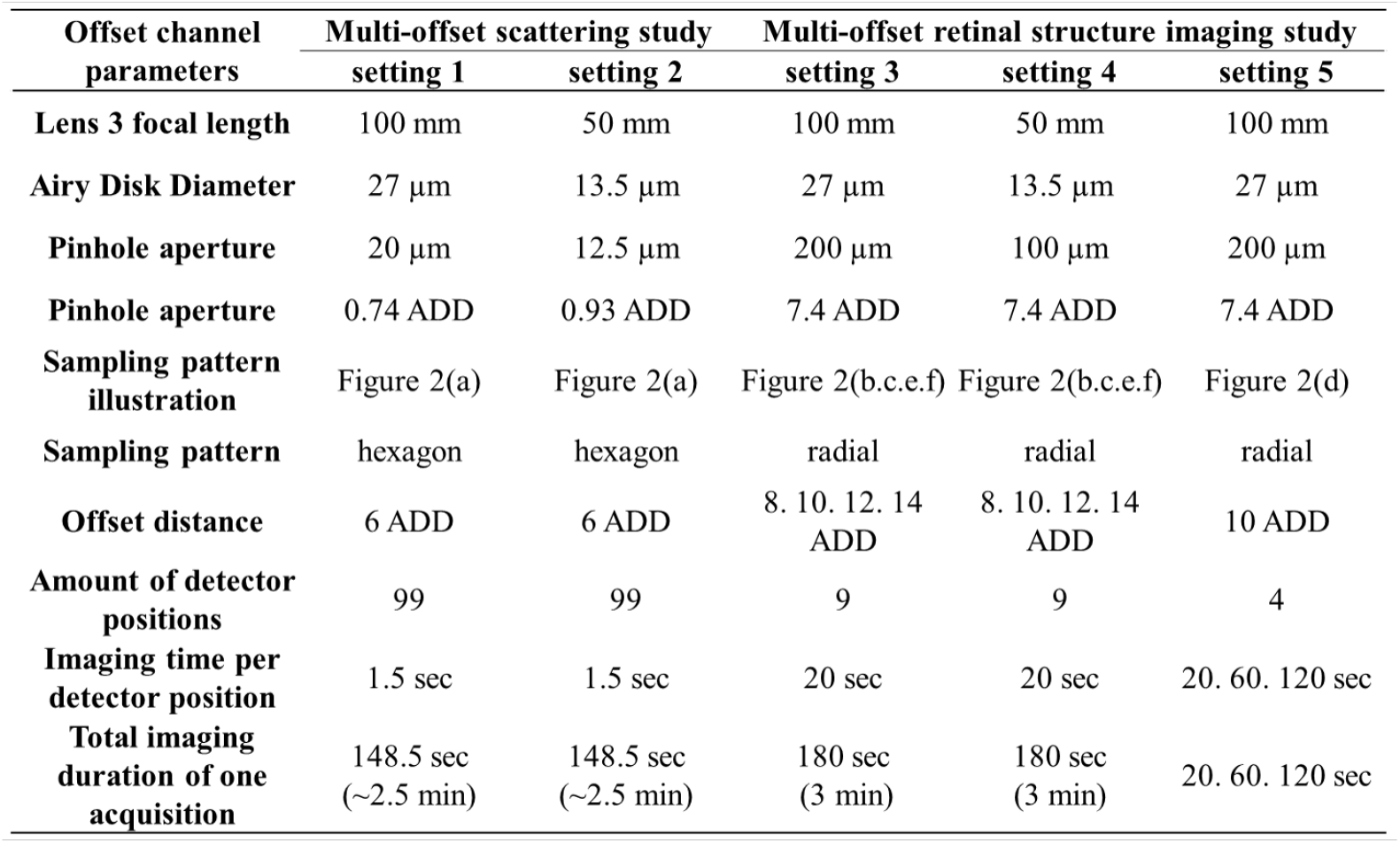
Offset channel parameters

### 2.2. Study design and Imaging protocol

We started by evaluating the optimal multi-offset configuration on one participant (participant 1) through the two following studies. We first focused on one specific retinal eccentricity (roughly 9° temporal (T), 3° superior (S)). The 1.5° x 1.5° field of view (FOV) was kept constant throughout image acquisition.

#### Multi-offset scattering study

Offset detection settings 1 and 2 are used for studying the multi-offset scattering properties (Table 1). We used a small pinhole aperture (0.74 and 0.93 ADD respectively) and the offset detector followed a hexagonal pattern with a lateral spacing of 6 ADD between adjacent detection positions in each row (see Fig. 1 (b)). The maximum offset distance from the center is 27 ADD in all directions, resulting in a total of 99 positions.

#### Multi-offset retinal structural imaging study

A larger pinhole aperture is used (7.4 ADD) to increase the signal at the detector for formation of the offset images and the offset detector is sequentially positioned in a circular pattern with a total of 9 positions (including the center position). To study the optimal offset detector position pattern for imaging retinal structures, particularly the ganglion cell layer, we first carried out several imaging sessions with the offset detector placed at four offset distances, 8, 10, 12, 14 ADD (settings 3 and 4 from Table 1 and Figure 1 (c-f). We started at 8 ADD as closer offset pinhole distances empirically showed residual confocal signals reaching the multi-offset detector. We also evaluated the effect of exposure duration for a single offset position. To improve image quality and to study the effect of exposure duration on the signal-to-noise ratio (SNR), we carried out a series of imaging session with longer exposure in 2 pairs of diagonal offset positions. At each position, we acquired 6 sets of 20-second videos, 2 sets of 60-second videos, and a 120-second video.

#### Validation of offset pattern distance on several participants

With the 8 or 10 ADD offset distance pattern (Fig. 1 (c) and (d)), we acquired several image sequences at different retinal locations in multiple participants. Retinal eccentricities imaged can be categorized into three regions: (1) temporal to the fovea within the macula: 4-9.5° temporal and 0-1.5° superior; (2) peripheral retina: 10-14° temporal and 3° both superior and inferior; (3) nasal superior to the fovea: 6-10.5° nasal and 8-9° superior (Fig.5). At each retinal location, offset images were acquired at 9 positions sequentially (Fig. 1 (c)) and a confocal video was simultaneously recorded. The ability to achieve images of the RGC layer was very sensitive to the focus of the AOSLO. We selected the retinal layer for the acquisition by changing the defocus on the deformable mirror and using the confocal channel as a guide to determine the deepest depth where the nerve fiber bundles were still visible in the confocal image. Figures 1 (d) and (e) show examples of averaged images generated from the confocal channel and one offset channel respectively after co-registration. Image sequences were acquired with an exposure time of 20s per aperture position for most experiments.

### 2.3. Image processing

All image processing was done using Matlab (R2019b, The MathWorks, Inc., Natick, MA)).

#### Image sequence registration

The sinusoidal artifact in the image sequence was corrected using previously published methods [11]. Images were then registered using a custom-developed registration algorithm developed in-house and described in detail elsewhere [9]. Offset aperture images were co-registered using the confocal sequence as the motion reference. The same reference frame was used for all confocal channel image sequences to ensure that the structure imposed by the confocal channel did not change at all between the different offset aperture images.

#### Image combination

From the registered sequence we generated average images for each offset position (hereafter called ‘offset images’). These offset images were then linearly combined to enhance the phase contrast using the following relationship:

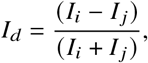

where *I_d_* is the obtained difference image (hereafter called ‘difference images’) and *I_i_* and *I_j_* are two offset images generated from two opposite pinhole positions. It should be noted that the background contribution of the resulting difference image is only partially mitigated, as some low frequencies remain.

#### Smart filter method

To enhance even more the signal of interest compared to the background, we filtered and combined images using a previously presented spatial frequency-based image reconstruction method (SMART filter), which was fully detailed elsewhere [7]. The SMART algorithm used here to generate the multi-offset images allows us to combine all difference images from the four directions.

To further mitigate the contribution of the background, a high pass filter with a normalized cut-off frequency of 0.02 was applied. Finally, we enhanced the local contrast of the images by applying contrast-limited adaptive histogram equalization (CLAHE) using the built-in Matlab function with 0.004 as the contrast enhancement limit.

### 2.4. Data Analysis

Images were scaled using the Gullstrand simplified relaxed eye model to estimate the value of microns per degree depending on the axial length of the participant.

#### Computing the Power Spectral Density

We computed the Power Spectral Density (PSD) to evaluate the distribution of power within the spatial frequencies corresponding to the cell mosaics under investigation. Using Matlab, a 2D Fourier transform was applied to the image. Then, we computed the square of the modulus of the Fourier transform. A radial average is then generated and divided by the value of the zero frequency to display the normalized PSD over the spatial frequencies (in cycles per mm). To enhance the energy peak at the spatial frequencies of the ganglion cells we fitted an exponential curve to the data and subtracted it from the initial data. This residual PSD curve displays a distinctive energy peak at a spatial frequency of the spacing of the detected cell mosaic.

#### Computing the Michelson Contrast

The Michelson contrast metric was computed on multioffset images obtained from the radial patterns with the four offset distances {8, 10, 12, 14} ADD and for each ADD size 13.5 μm and 27 μm. For each image the Michelson contrast value of 26 regions including a ganglion cell (white rectangles in Fig.2 (e)) was averaged and the standard deviation was computed.

**Fig. 2.**
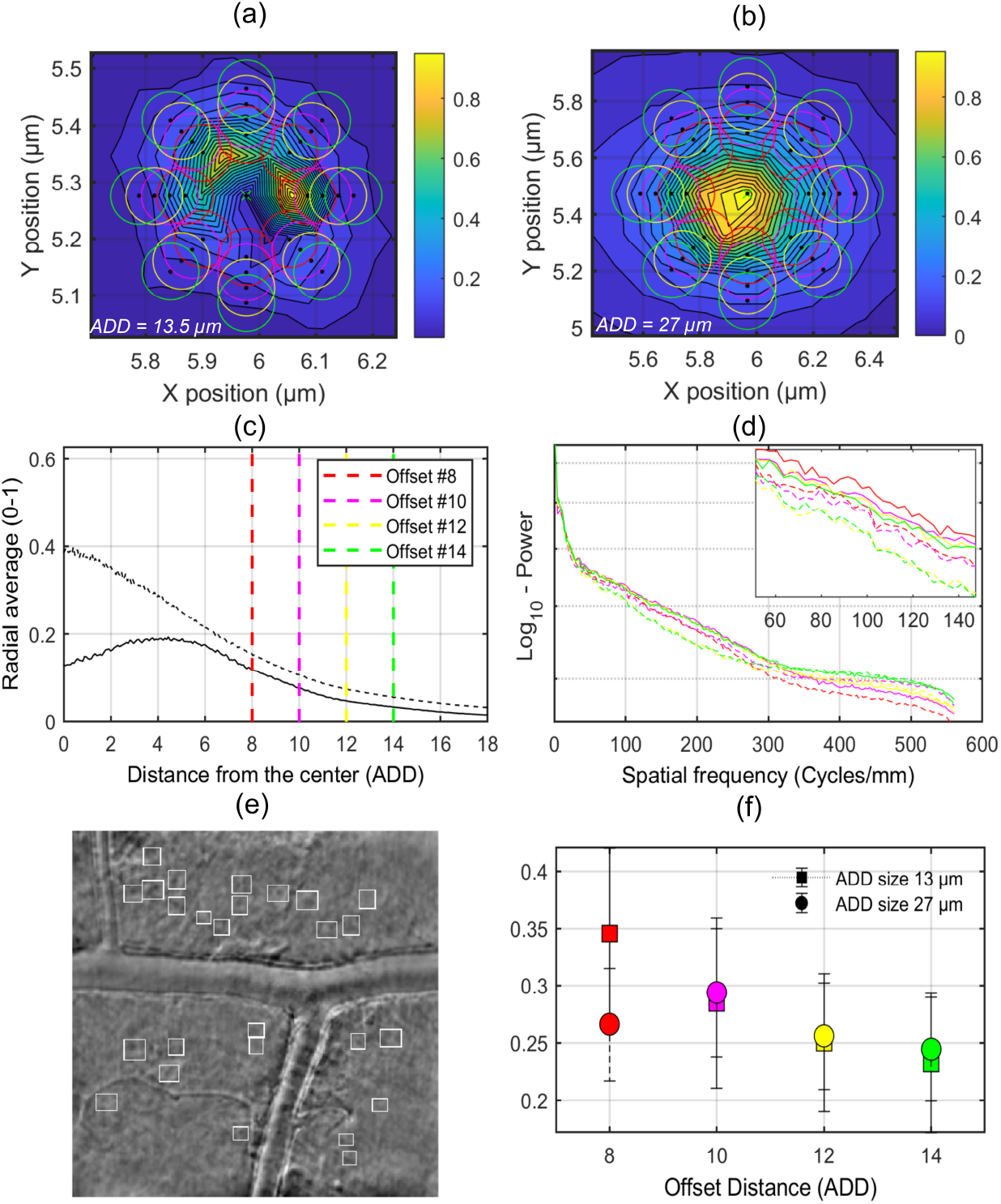
Image quality metrics. Metrics computed on images of participant 1 at 8° eccentricity. Metrics represented in {red, magenta, yellow, green} were calculated on images acquired at offset distances {8, 10, 12, 14} ADD respectively: (a,b) Surface plots of the pixel intensity at the xy multi-offset detection plane for ADD 13.5 μm and 27 μm respectively; (c) Radial averages of the pixel intensities with respect to the offset distance at the xy multi-offset detection plane. Solid and dashed lines represent 13.5 μm and 27 μm respectively; (d) Power spectra of averages from acquisitions with the four offset distances (8-14 ADD), a zoom highlights the increase in energy around the spatial frequencies corresponding to the ganglion cell spacing. Solid and dashed lines represent 13.5 μm and 27 μm respectively; (e) Multi-offset image showing the region of interest evaluated, acquired with an offset of 8 ADD and 13.5 μm ADD. White rectangle identify ganglion cells were the Michelson contrast was computed; (f) Average Michelson contrasts computed on a range of cells marked with a white rectangle on (e) for multi-offset images obtained with (8-14 ADD) offset distances and with both 13.5 μm and 27 μm ADDs.

The Michelson contrast of the same profile segments of some retinal structures (inner segment, ganglion cell, vessel wall) was computed on the individual offsets, the difference images of opposite offsets and multi-offset images of the retinal region shown in Fig.3.

**Fig. 3.**
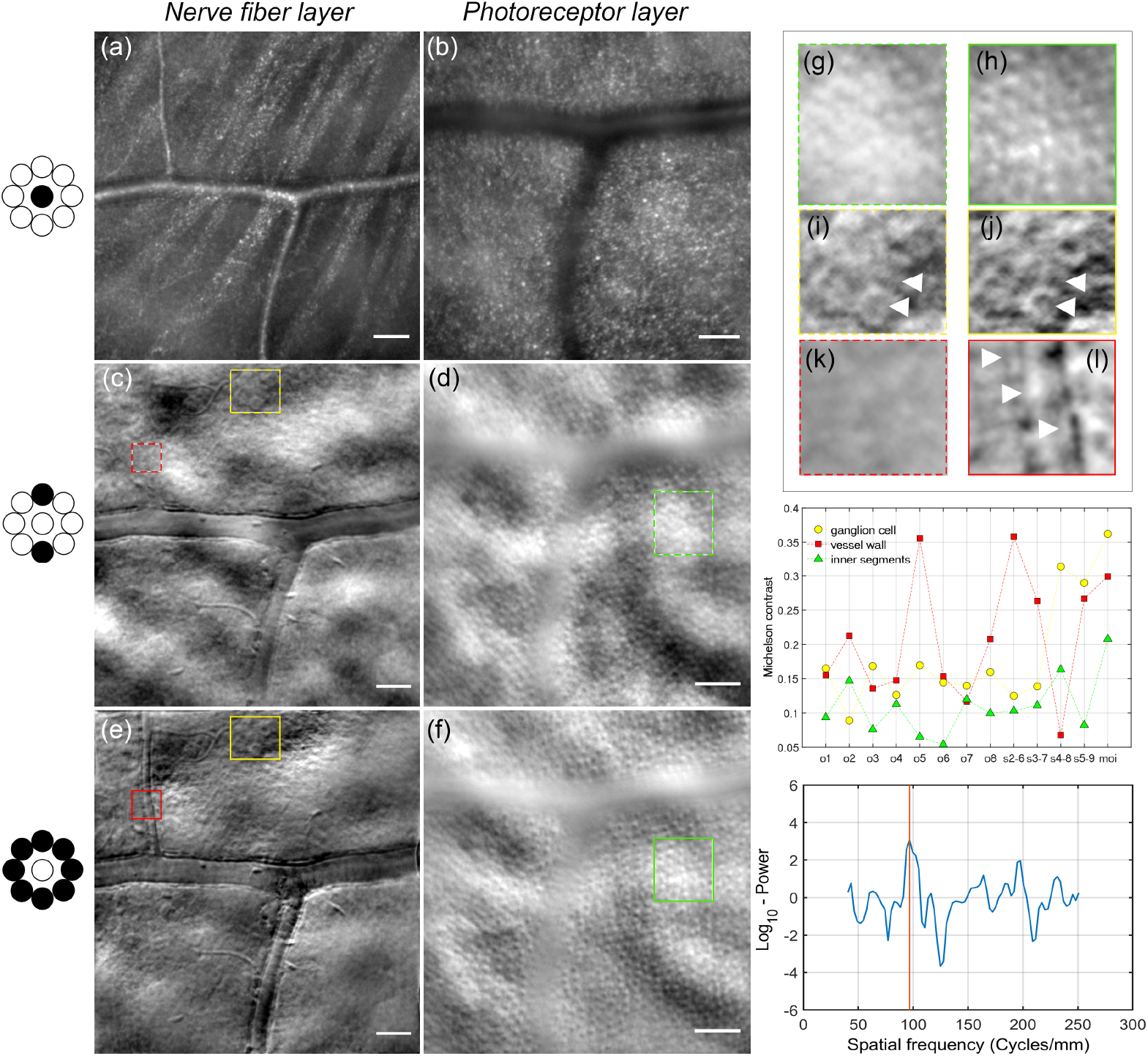
Multi-offset images. All images are from participant 1 at 8° eccentricity. (a,b) Confocal images; (c,d) Difference images; (e,f) Multi-offset images; (g-l) Enlarged regions displayed in green, red and yellow rectangles with a dashed line in the difference images (c,d) and a solid line in the multi-offset images (e,f). White arrowheads show structures present in the multi-offset images such as details of the ganglion cells or the vessel wall but absent or low contrasted on the difference image. (m) Michelson contrast computed on the segments shown in from opposite offsets zoomed regions (g-l) for offset images {o1-o8}, difference images {s1-s4}, and the multi-offset images {moi}. (n) Logarithmic plot of the residual of the difference between the Power Spectrum Density of image (e) and its exponential fit. A peak is detected at a 92 cycles per mm (which corresponds to an average cell spacing of 12 μm). Scale bars are 50 μm.

### 2.5. Participants

For this study, 18 healthy participants were imaged (6 male and 12 females, ages ranging 20-64 yrs old). Individual participant characteristics including age, sex and eye that was imaged can be found in Table 2. Participants were free from eye conditions including congenital eye diseases, recent ocular infection, glaucoma, and recent eye surgery. They also had clear lenses, deep and quiet anterior chambers, and a refractive error no more than –4D. Written informed consent was obtained from all participants following an explanation of experimental procedures and risks both verbally and in writing. All experiments were approved by the University of Pittsburgh Institutional Review Board and adhered to the tenets of the Declaration of Helsinki. The imaging powers used were 590 μW for the 795 nm center wavelength SLD and 21 μW at 909 nm for the wavefront sensing beacon laser diode. All cumulative light levels for multi-wavelength exposures were below the limits specified by the ANSI [12]. Before each imaging session, we applied eye drops for pupil dilation and cycloplegia (one drop each of 1% Tropicamide and 2.5% Phenylephrine Hydrochloride).

**Table 2.** Participants information

|  | Category | Participants (n=18) |
| --- | --- | --- |
| Sex | Male | 6 (33.3%) |
|  | Female | 12 (66.7%) |
| Eye | OD | 10 (55.6%) |
|  | OS | 8 (44.4%) |
| Age (years) | | 30 ± 10 |

## 3. Results

### 3.1. Comparing 4 detection configuration setups

Using the radial detection configuration with 8 offset positions we evaluated four offset distances on participant 1 ({8, 10, 12, 14} ADD) as shown in Fig.1 (c-f) in terms of amount of signal detected, power spectrum density at spatial frequencies corresponding to ganglion cells spacing and Michelson contrast of detected cells. We show in Fig. 2 that offset distances 8 ADD and 10 ADD lead to higher signal and image quality for both ADD sizes 27 μm and 13 μm at the multioffset plane (settings 3 and 4 in Table 1).

#### Multi-offset scattering quantification

The average intensity on the xy plane at the multi-offset detector plane is plotted in Fig.2 (a) for ADD = 13.5 μm and (b) for ADD = 27 μm of the field of view shown in (e). A colormap shows the distribution of the multiply scattered signal around the center at the detector plane after the confocal part of the beam has been removed by the reflective pinhole. The four multi-offset detection patterns described in Fig.1 (c-f) have been superimposed to the intensity plots to show their respective position with respect to the intensity distribution of the multiply scattered signal. In Fig.2 (c) the radial average of (a,b) are shown with respect to the offset distance in ADD. Dashed lines indicate the position of the offsets for the four offset distances evaluated showing how at 12 ADD offset distance the amount of signal is already half what it was at 8 ADD.

#### Multi-offset imaging quality quantification

For each offset distance we computed the PSD (Fig.2 (d)) which shows an increase in energy at the spatial frequencies corresponding to the ganglion cell mosaic spacing (around 80-100 cycles/mm) for the lowest offset distances (8,10 ADD) represented in red and magenta respectively. We also noticed a peak increase of the PSDs corresponding to an ADD size of 13.5 μm represented by solid lines. On the contrary the average Michelson contrast values computed on cells indicated in Fig.2(e) with white rectangles seem to match for both ADD sizes, with a slight difference for the 8 ADD distance. Nevertheless, similarly to the previous metric, 8-10 ADD offsets lead to more highly contrasted profiles (with well defined cell borders) and higher Michelson contrast.

### 3.2. Detection of Ganglion cells

Figure 3(e) illustrates an example of ganglion cells obtained with phase contrast through multi-offset imaging using a radial pattern with an offset distance of 8 ADD (with an ADD size of 13 μm) on participant 1 at 8° temporal. A zoom indicated in Fig. 3 (i-j) show the details of a cluster of ganglion cells. A peak of the Power Spectrum Density of the multi-offset image (e) was detected at 92 cycles/mm (Fig.3 (n)), which corresponds to a cell spacing of 12 μm, in accordance to the average size of the ganglions. In Fig.3 (m) we show how the phase contrast of the ganglion cells increase by computing the difference images between opposite offsets, and even further when computing the multi-offset image (or moi in the plot). This improvement in contrast can be appreciated in the zoom (j) compared to (i).

### 3.3. Multi-layer imaging of the retina with the radial pattern

The multi-offset AOSLO technique enables us to generate high resolution images of distinctive layers of the retina as illustrated in Fig. 3. With the confocal detector, highly reflective layers can be imaged, resolving fiber bundles (Fig.3a) as well as individual photoreceptors (Fig.3b). With the multi-offset pattern set to 8-10 ADD we evaluated the contrast enhancement of this radial pattern compared to single offsets and difference images for other structures of the retina. Figure 3 (h) show the resolution of inner segments in the multi-offset image unidentifiable on the difference image Fig.3 (g). This is further confirmed by the Michelson contrast metric in Fig.3(m). Finally, some retinal structures such as vessels display a very directional scattering of the light. This leads to the absence of multiply-scattered photons from vessels reaching some offset pinholes therefore unable to resolve them as shown in the difference image Fig.3 (c). The vessel wall clearly identifiable in multi-offset image (l) by arrowheads is absent for the direction of the difference image (k).

### 3.4. Reproducibility

We demonstrated the repeatability of the technique in Fig.4 where the same ganglion cells can be observed over different time points. We identified examples of the same cells (indicated with arrowheads) on images acquired with a difference of weeks (a,b) as well as on images acquired with a difference of minutes (b,c,d). A zoomed region shows the details of one of these cells.

**Fig. 4.**
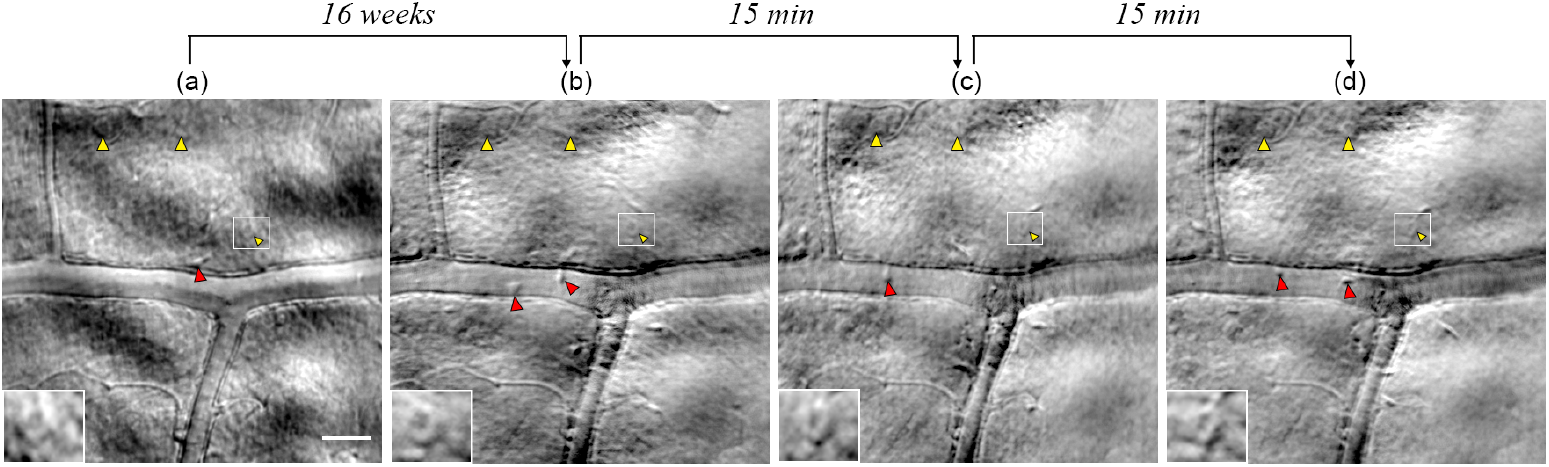
Reproducibility. Images acquired at 4 different time points showing that the same cells can be imaged over time. yellow arrowheads indicate examples of ganglion cells visible at the different time points. Red arrowheads identify putative microglial cells. Scale bar is 50 μm.

We also noticed when imaging the same region over time other structures in this layer that were moving over a few tens of microns. We hypothesised by the size and motion that these could be microglial cells that seem to move over a timescale of several minutes. In Fig.4 we identified these cells with white arrowheads. The motion can be better observed in Visualization 1 where the arrowheads follow the motion of putative microglial cells.

### 3.5. Validation on several participants

We validated our radial multi-offset pattern with an offset distance of (8-10) ADD on other participants by detecting ganglion cells. Figure 5 show images of RGCs revealed on participants 1-7 at different eccentricities on multi-offset images. As mentioned in previous studies [3] the polarity of cells imaged with phase contrast imaging in the retina can be positive (blue curve in Fig.5 (n)) or negative (black curve in Fig.5 (n)). We identified some of the ganglion cells with white and yellow arrowheads for cells with positive and negative polarity respectively. Cells in positive polarity reach its maximum intensity value of the profile at the center while those with negative polarity reach their maximum at the wall and minimum at the center. Inverting the image allows to move from one polarity to another. As shown in several participants in Fig.5 cells with different polarities are simultaneously captured in the same region with our multi-offset technique. We also observe a range of soma sizes, with cells as small as 6-8 μm going up to 20 μm. A table in Fig.6b summarizes the averages computed on several participants of soma sizes, spatial frequencies of the PSD peaks shown in Fig.6a and the density and cells spacing derived from these spatial frequencies.

**Fig. 5.**
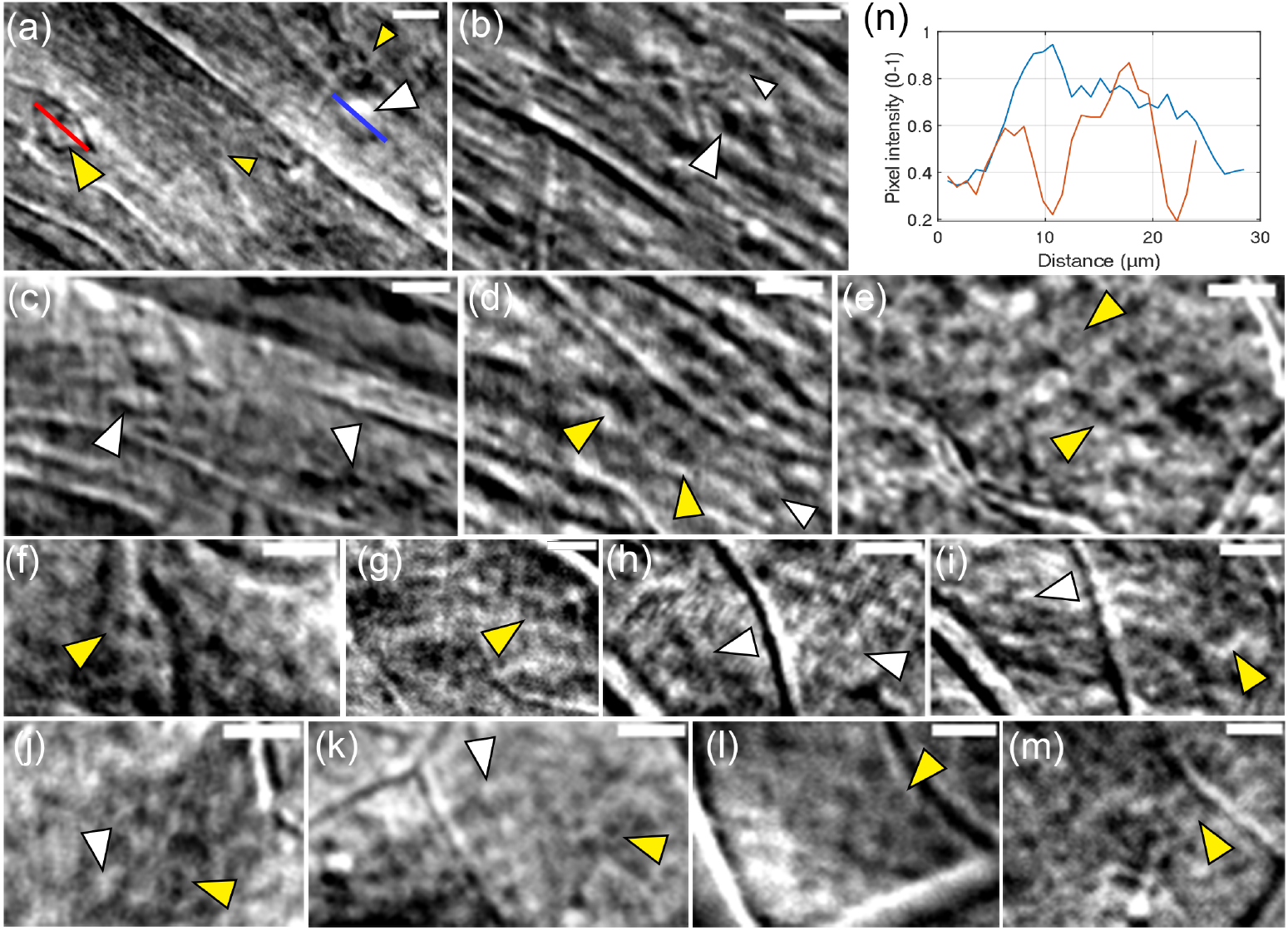
Validation on other participants. Multi-offset images on participants 2 (a-e), 3 (f,j), 4 (g), 5 (h,i), 6 (k,m) and 7 (l) on different eccentricities (around 12° Nasal (a-d), around 6° Temporal (e-i) and around 8-10° Temporal (j-m)); (n) Intensity profiles from red and blue segments shown in (a) of two cells with inverted polarities (red for negative where the center is dark and borders are brighter, blue for positive where the center is bright). White arrowheads identify ganglions with positive polarity while yellow arrowheads identify ganglion cells with negative polarities. Scale bars are 10 μm.

**Fig. 6.**
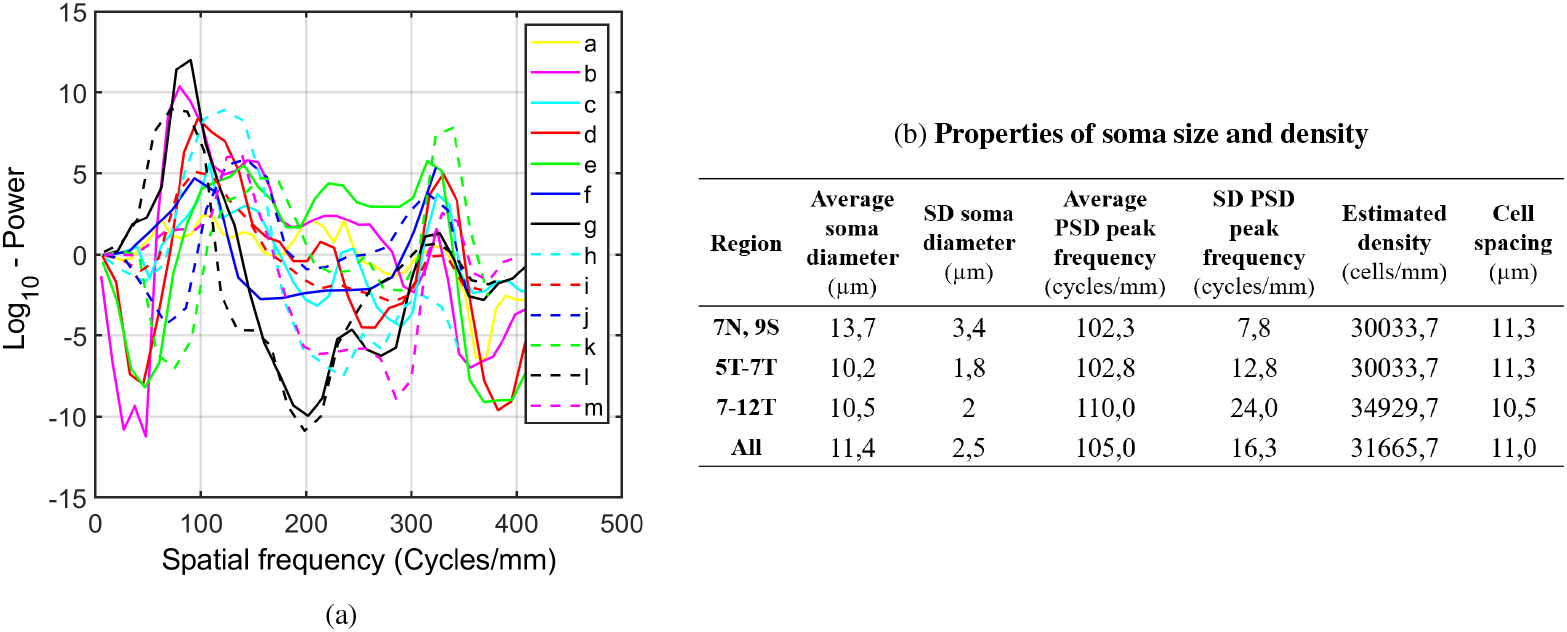
(a) Residual Power Spectrum Density plots of multi-offset images from Fig.5; (b) Average soma size, PSD peak spatial frequency, density and cell spacing for eccentricity ranges (7N,9S), (5T-7T) and (7T-1T) computed on participants (1-7).

## 4. Discussion

We implemented a new radial multi-offset detection AOSLO setup optimized to enhanced the phase contrast signal from RGCs. Additionally, this new optical setup has obviated some of the major limitations of our previous approach [8] and demonstrated robust, repeatable imaging of RGC layer neurons using multi-offset imaging, with relatively high-contrast in humans for the first time. This builds on our previous work and demonstrates that similar contrast levels to what we had shown previously only for anesthetized monkeys at high light levels can be obtained in humans at safe light levels with relatively short duration exposures, on the order of 10 to 20 seconds per aperture position.

One of our main aims here was to overcome previous experimental limitations to improve the phase contrast signal of the ganglion cell layer in humans *in vivo* in AOSLO. By splitting the confocal and multi-offset detection channels with the custom-made reflective pinhole we were able to use the same imaging wavelength for both channels, in a similar way to nonconfocal split-detection [6]. This was a major improvement as it eliminated any possible blur induced from transverse chromatic aberration when using a different wavelength for confocal imaging and for tracking eye motion. We usually assume that there is a fixed Transverse Chromatic Aberration (TCA) between each imaging channel but changes in pupil position can change the TCA and this can be rendered as motion blur when assuming a fixed offset. This is not an issue when we use the same wavelength of light to track the eye as we do for the multi-offset imaging. The use of a single near infrared imaging source permitted the use of higher light levels than for our previous multi-wavelength implementation of multi-offset imaging that used a combination of visible and near-infrared (NIR) light. Removing the hazards associated with visible light exposures permitted us to use more NIR light for imaging safely.

The principle behind this technique, previously described [13, 14], is to detect the phase signal of the photons after going through the cells by converting it into intensity at the detection plane. It is achieved by shifting the detection pinhole away from the center in different directions. The light going through the cells is deviated differently depending on the refractive indices and therefore photons from different regions of the cell do not reach the same offset aperture leading to differences between adjacent tissues with different refractive indexes in transparent cells [3]. In Visualization 2 we show a Fourier Transform of the same region shown in Fig.3 (a) acquired with each of the 8 offsets showing the spectral information rotating around the center as we move through the pinholes. As the contrast of the offset images results from the differences in phase of the detected photons, this variation in spectral content is directly linked to a variation in the detected phase. In addition to observing this variation in spectral information when moving the pinhole around the center we explored the distribution of multiply scattered light, which carries the phase information, in the detection plane. We noticed that although there is some directionality in the signal distribution (Fig.2 (a,b) and Suppl.Fig.1) these photons are mostly scattered all around the center. Furthermore, the fact that the scattering directionality is most likely related to type and orientation of retinal features imaged such as vessels or nerve fibers, a radial pattern enables to capture photons deviated in all directions without the need to determine first their main direction of scattering.

In our current technique, which was inspired by phase contrast microscopy [15, 16] and from previous off-axis configurations in the ophthalmic community [5, 6, 17, 18], we generated what we call difference images by subtracting the images from the opposite offsets to cancel the background as well as absorption signals leaving only the phase signal. The highest phase contrast is theoretically generated by subtracting opposite pinholes, as with homogenous illumination they should lead to opposite phase values, and therefore double the signal when subtracted. However, as our background was not homogenous, we were unable to cancel it fully as can be seen in Fig.3(c) where the cell signal remained modulated by the low frequencies of what we hypothesize corresponds to a choroidal signal. This is likely due to the apparent displacement of the low spatial frequency features of the background with change in aperture position (this apparent parallax in the images causes the background to appear to move when the aperture is offset to different positions).

Given the weak phase signal coming from the ganglion cells we explored different parameters of this radial configuration, such as the offset distances and the size of the ADD at the detector plane, that would increase our image quality metrics. We had two objectives, firstly to evaluate if the relative distance in ADD would remain constant by changing the magnification of the detection plane, and secondly to determine the effect of a lower magnification (ie a smaller ADD size at the detection plane) on the phase contrast. The starting offset distance of 8 ADD was selected by placing the pinhole at a distance where the residual confocal (singly scattered light) signal was not visible on larger vessels on the multioffset detection channel. The sampling of the multiply scattered light also allowed us to evaluate the signal decrease with respect to the distance from the center of the Airy disk (Fig. 2c), which displays a similar decay for all the tested regions and ADD sizes (see Suppl.Fig.1). We selected a range of 8-10 ADD offset distances which captured the largest amount of the multiply scattered signal to have highest SNR.

The results in Fig. 2 (d,f) showed for both ADD sizes respectively that an offset distance of 8-10 ADD provided the highest contrast. This is most likely governed partially by the scattering distance that leads the signal to reach higher intensity at these distances (Fig. 2(a,b)) and related to the difference in refractive index between ganglion cells and medium. When the latter increases the difference between the pixel intensity of the cell border with respect to the medium also increases as shown in Fig. 2 (f). The fact that this optimal distance seems to remain constant in ADD would facilitate the implementation of this setup on other AOSLO systems. It also implies that by simply changing the ADD size at the detector plane, different fiber optic bundles could be used to achieve this configuration. Nevertheless it is worth mentioning the slight enhancement of the energy peak observed at the spatial frequencies of the ganglion cells mosaic when decreasing the Airy disk diameter (see Fig. 2(d)). This suggests that other groups might benefit from decreasing the magnification of the offset detection path lens telescope in their AOSLO that is usually set to a magnification that facilitates alignment with larger Airy disk diameters.

We also verified the ability to reproduce the same cell over time by recording several acquisitions with the same detector configuration and eccentricity at different time points. The presence of the same cellular structure within longer and short intervals demonstrated the monitoring capacity of this technique. Moreover, in Visualization 1 we can see how ganglion cells remains still in the background while other structures navigate through this layer over intervals of a few minutes.

We validated our detection scheme on several participants as shown in Fig.5. We show for several multi-offset images from three different eccentricities cluster of ganglion cells with sufficient contrast to compute the properties of soma size and cell spacing presented in the Table in Fig. 6 (b). Not all participants displayed the same contrast levels, which can be contributed to common issues such as non-optimal tear film layer, watering eyes or excessive fixational eye movements. Despite these non-optimal imaging conditions we were able to detect ganglion cells on a range of participants and image the same cells over time.

Another technology targeting these cells, described by Liu et al. [19], is adaptive optics optical coherence tomography (AO-OCT). This technique detects singly back-scattered light from transparent retinal tissue and is thought to exploit organelle motility inside ganglion cell somas to generate high contrast images of RGCs, which allowed them to identify ganglion cell somas and quantify structural parameters such as soma density and diameter. Although our cell contrast is lower compared to AO-OCT, radial multi-offset detection is the only other imaging technique capable of detecting human ganglion cells *in vivo*. Unlike AO-OCT, the contrast does not originate in organelle motility but is a result of the difference in refractive indices n between the tissues. This implies that although revealing the same cells, the two techniques detect different cellular structures, potentially providing complementary information about the RGC layer. This contention is supported by our previous work in monkeys that showed that multi-offset imaging may potentially reveal sub-cellular structures, such as the cell nuclei [8], whereas these features are not visible in the AO-OCT images of RGC layer neurons [19]. Future development could aim to implement an AOSLO/AO-OCT setup similar to the work of Dubra’s group [20] to overlay both signals from ganglion cells.

We hypothesized that we are able to detect microglial cells (see Visualization 1), which would allow us to monitor their dynamics similarly to what AO-OCT groups have been able to achieve [21]. However, given that AO-OCT imaging requires acquiring several data volumes and averaging images over a significant amount of time, our multi-offset technique may reach a higher temporal frequency and therefore finer motion monitoring of these cells.

One of the caveats of sequential acquisition is that it increases the total acquisition time, which again is not ideal from the patient imaging perspective. Parallelizing the offset imaging fiber bundle and recording each channel simultaneously would speed up imaging as light waste would be significantly reduced. We are currently implementing a fiber bundle using the parameters from the optimized radial pattern designed in this study to decrease the exposure time and more importantly to guarantee simultaneous acquisition of the multiply scattered light by all offset pinholes as fixational eye movements currently lead to slight displacements with respect to the different pinholes during the acquisition. This motion probably leads to slight changes in the expected detected phase eventually affecting the difference images. With the fiber bundle we would only need several seconds of exposure time enabling us to follow the motion of these putative microglia at a very fine temporal scale or achieve imaging of participants or patients with poorer fixation.

Finally we demonstrated not only the capacity of our multi-offset radial pattern to detect other retinal structures typically imaged with phase contrast (i.e.vessel walls, capillaries or inner segments), but also showed through the image quality metrics an enhancement in contrast by combining the radially distributed pinholes. This pattern enables us to mitigate the effect already described in the literature [5] of disappearing vessels depending on the relative position of the offset pinholes due to the directionality of the scatter. This showed that for offset aperture imaging of retinal vasculature, a pattern that covers all angles is necessary to capture a full perfusion map of all the vessels (see Suppl. Fig. 2).

## 5. Conclusion

We optimized the multi-offset imaging using an improved single-wavelength optical system to mitigate the impact of the limitations imposed by our previous multi-wavelength approach. We implemented a radial multi-offset detection pattern with which we achieve repeatable high contrast images of retinal ganglion cells at different eccentricities and participants. We demonstrated the enhancement in contrast with respect to more standard phase contrast imaging techniques such as offset or split-detection, and with respect to previous multi-offset setups. Multi-offset imaging could be a complementary tool to AO-OCT, for retinal ganglion cell detection at cellular resolution, and even become a valuable tool for studying fast dynamics of microglia, and potentially other weakly scattering structures, in humans *in vivo*.

## Funding

This research was supported by a National Glaucoma Research Award from the BrightFocus Foundation (G2017082) to Ethan A. Rossi. It was also partially supported by a grant from Foundation Fighting Blindness (PPA-081900772-INSERM) and by departmental startup funds from the University of Pittsburgh to Ethan A. Rossi. This work was also supported by NIH CORE Grant P30 EY08098 to the University of Pittsburgh Department of Ophthalmology, the Eye and Ear Foundation of Pittsburgh, the NVIDIA GPU Grant Program, and from an unrestricted grant from Research to Prevent Blindness, New York, N.Y. USA.

## Acknowledgments

The authors would like to thank Austin Roorda for sharing his AOSLO software with us and Pavan Tiruveedhula for electronics fabrication, software, and technical assistance.

## Disclosures

Ethan A. Rossi has patents on some aspects of the technologies used to carry out these experiments (US10,772,496; US10,123,697; & US10,092,181).

## Notes

### Competing Interest Statement

The authors have declared no competing interest.

### Summary of Updates

We added more information about the design of detection pattern and more quantification analysis of the images. Supplemental files updated.

