## Supplementary material for "Design of a radial multi-offset detection pattern for *in vivo* phase contrast imaging of the retinal ganglion cells in humans": Suppl. Fig.1, Fig.2, Table 1

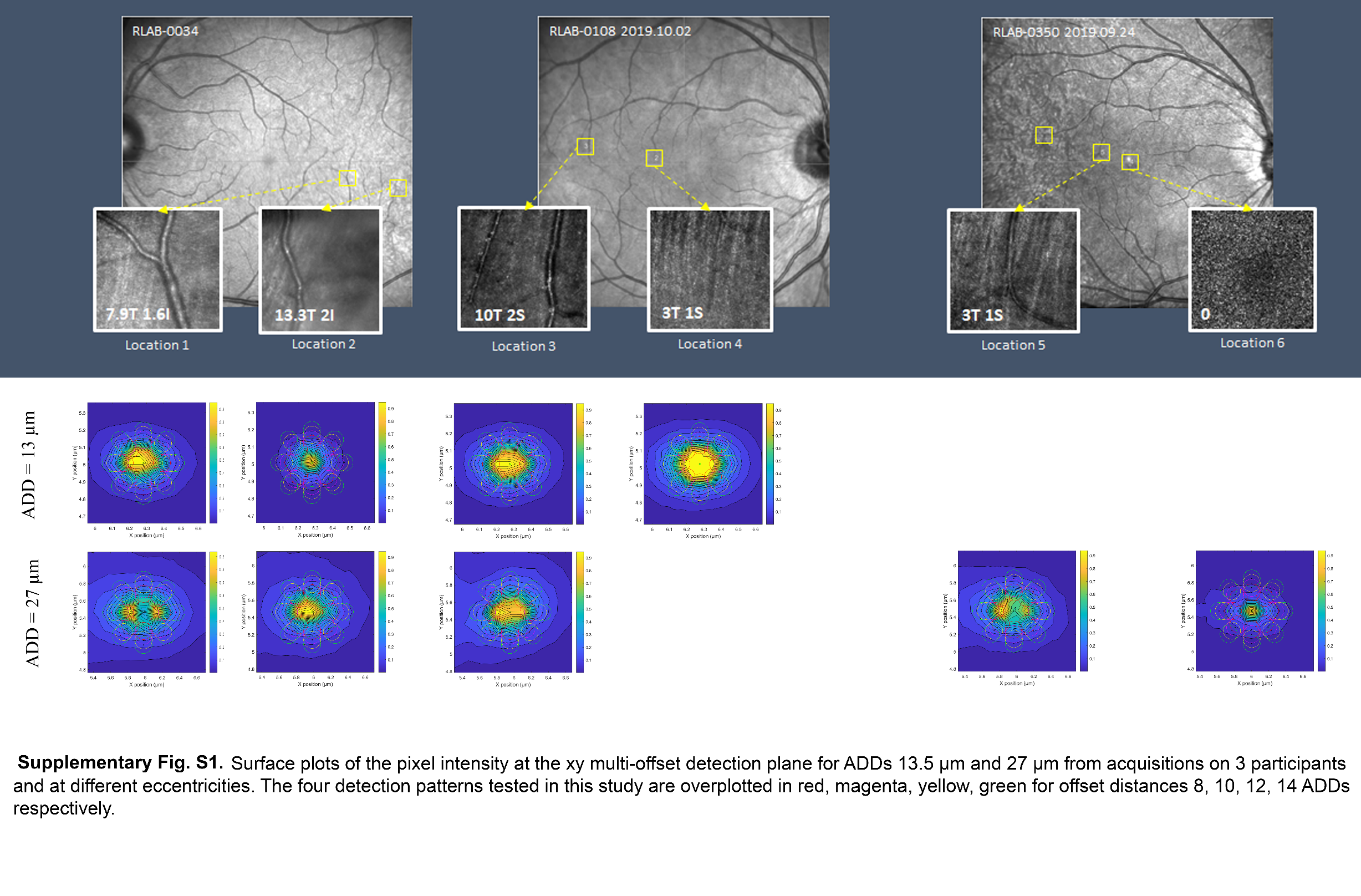

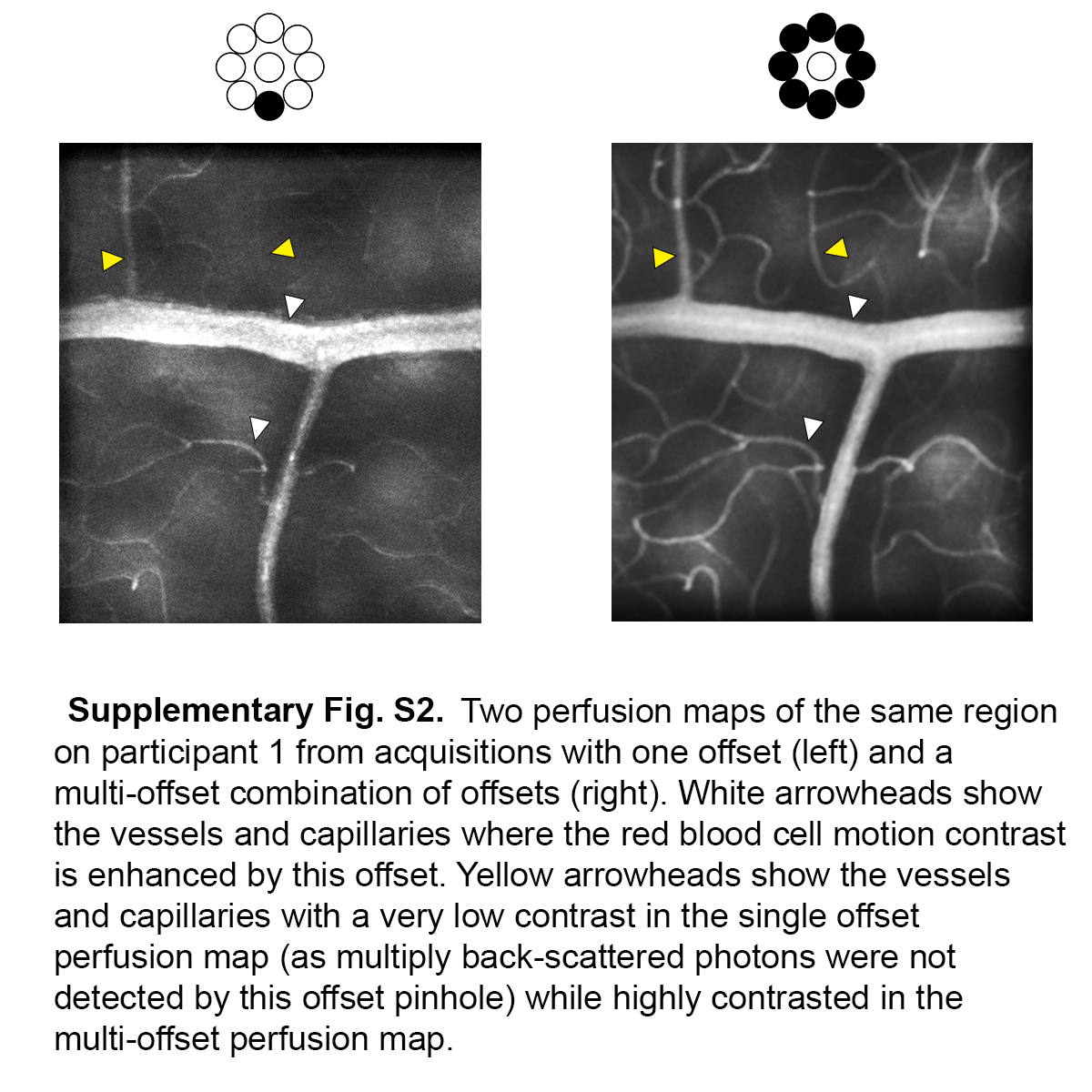

**Supplementary Table 1.** Comparison between the parameters of the previous and current multi-offset setups

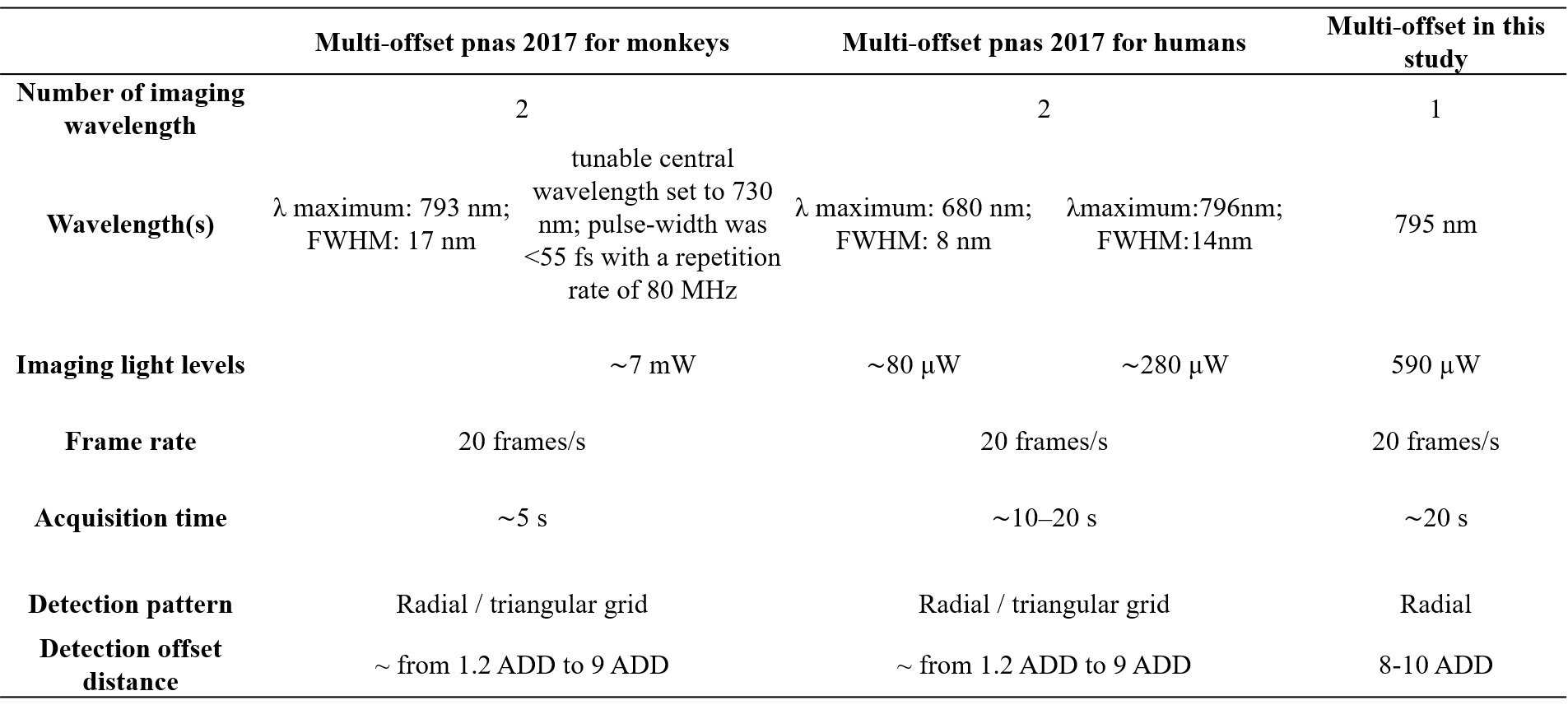
